## Supplementary material for "Oncogenic EFNA4 amplification promotes lung adenocarcinoma lymph node metastasis": Figure S1-6

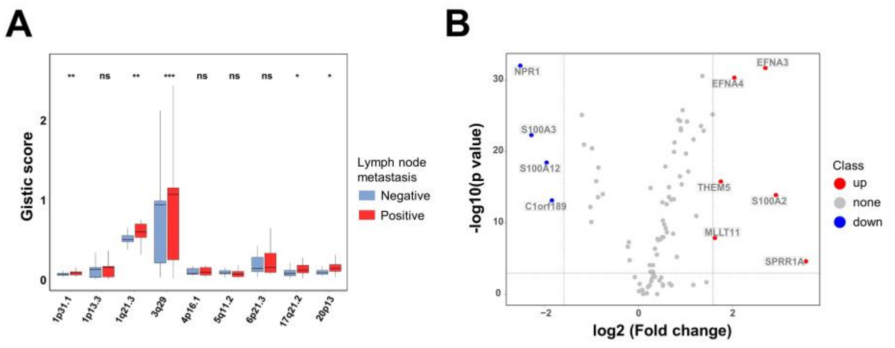


**Figure S1.** 1q21.3 amplification in metastatic lung cancer. (A) Gain of copy number and amplification of nine most common amplified cytobands in metastatic and nonmetastatic lung cancer. The box shows median value and 25th and 75th percentiles; (B) Volcano plot of the distribution of all differentially expressed genes with statistical significance and fold change at the 1q21.3 region between lung tumor samples and paired normal tissues. Significant p-values are represented as * $p < 0.05$, ** $p < 0.01$ and *** $p < 0.001$.

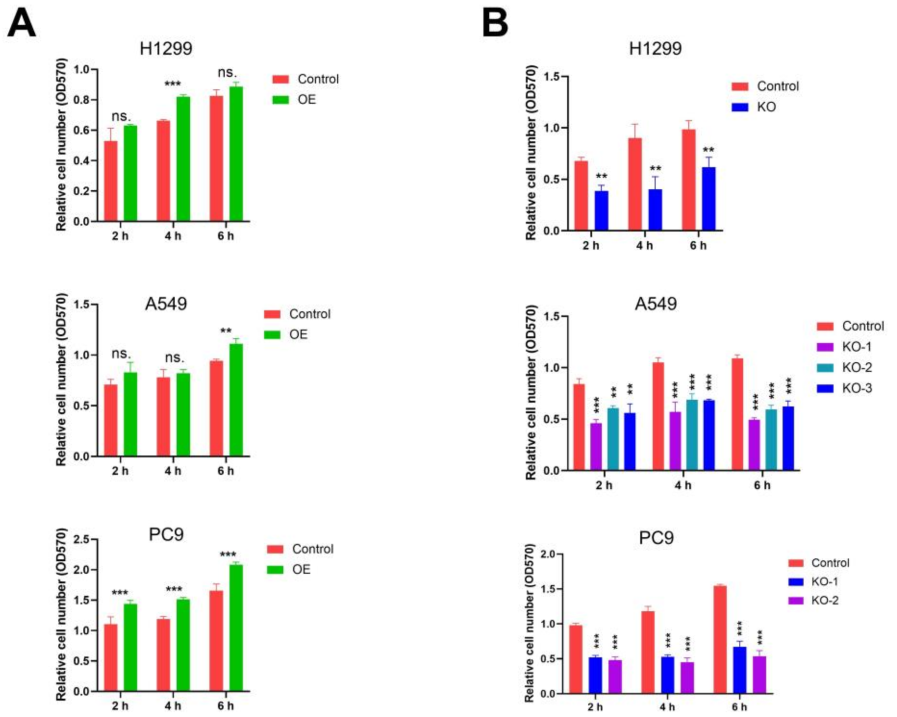


**Figure S2**. EFNA4 expression level is associated with cell adhesion. Cell adhesion assay. Adhesion ability of wild type, EFNA4-overexpressing (A) or EFNA4 deleted (B) cells was analyzed by counting cells using MTT assay. Data represented as mean ± SD of three independent experiments. OE: over expression; KO: knockout. Significant p-values are represented as * $p < 0.05$, ** $p < 0.01$ and *** $p < 0.001$.

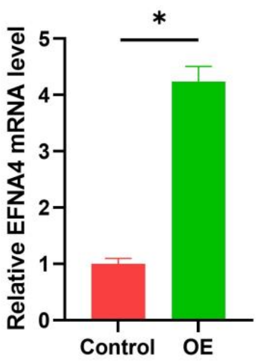


**Figure S3**. RT-qPCR was utilized to detect expression levels of *EFNA4* in tumor xenografts derived from *EFNA4* over-expressing or control A549 cells. Data are expressed as mean ± SD of three independent experiments. OE: overexpression. Significant p-values are represented as * $p < 0.05$, ** $p < 0.01$ and *** $p < 0.001$.

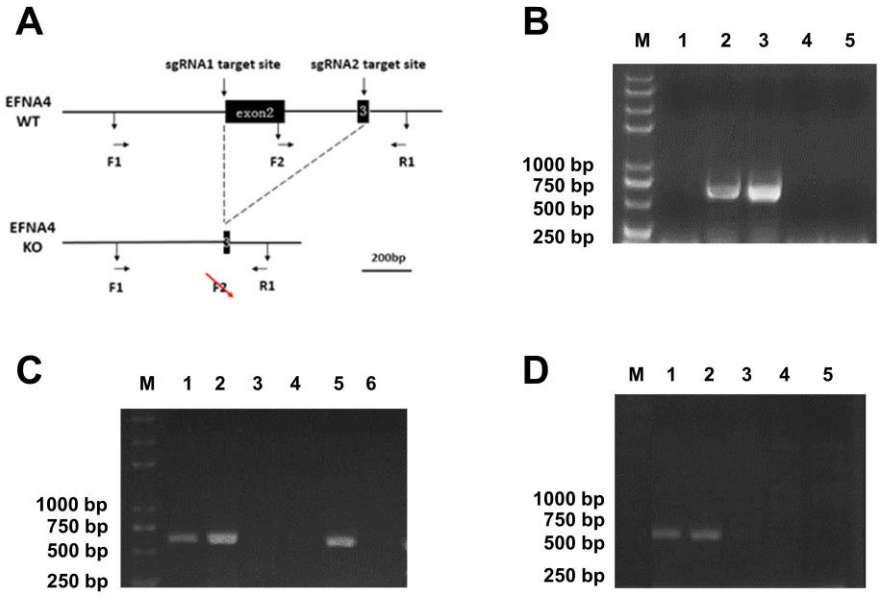


**Figure S4.** Knockout *EFNA4* in lung cancer cells using CRISPR/Cas9. (**A**) Schematic representation of the *EFNA4*-knockout model. F1/F2 represents upstream primer, R1 represents downstream primer; Agarose gel electrophoresis of the polymerase chain reaction (PCR) products of H1299 (**B**), A549 (**C**) and PC9 (**D**). Lane M shows DNA ladders. PCR products of homozygous knockout cell lines showed shorter length on the gel.

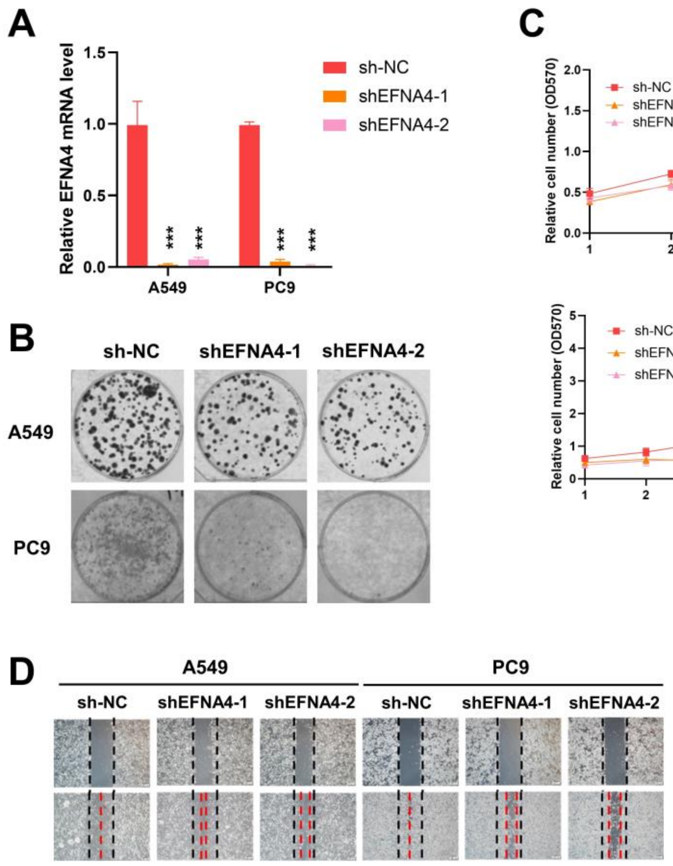


**Figure S5.** Downregulation of *EFNA4* inhibits the growth and migration of lung tumor cells. (**A**) qPCR data of EFNA4 mRNA expression in A549 and PC9 cells transduced with lentiviruses carrying vectors with the indicated shRNAs or empty vector (sh-NC). Relative *EFNA4* expression values were normalized to *ACTB* gene; (**B**) Representative images of methylene blue-stained colonies in A549 and PC9 cells transfected with EFNA4 shRNA or NC shRNA are shown; (**C**) MTT cell proliferation assay. A549 and PC9 control cells and *EFNA4* knockdown cells proliferation was measured by MTT method. Data are presented as means ± SD; (**D**) Cell migration analysis by wound-healing assay in *EFNA4* knockdown cells and control cells of A549 and PC9 cells. Data are expressed as mean ± SD of three independent experiments. Significant p-values are represented as * $p < 0.05$, ** $p < 0.01$ and *** $p < 0.001$.

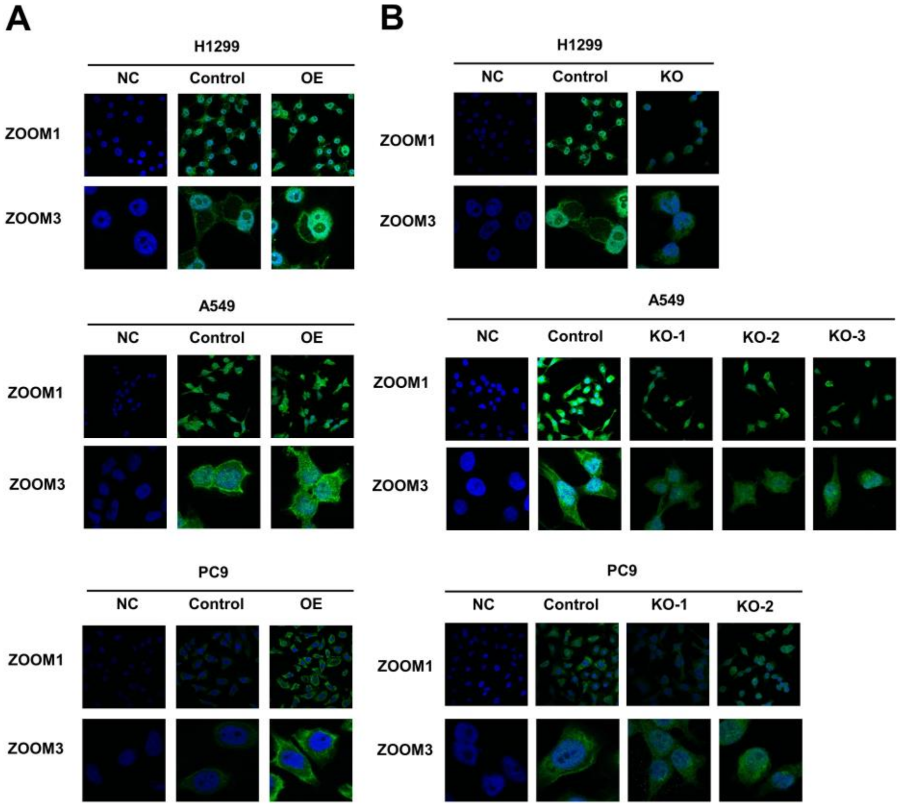


**Figure S6.** EFNA4 is located on the cell membrane in LUAD. (**A**) Confocal images of control cells and EFNA4-overexpressing cells labeled with anti-EFNA4 (green) and DAPI (blue), respectively; (**B**) Confocal images of control cells and EFNA4 knockout cells labeled with anti-EFNA4 antibody and DAPI, respectively. Panels show nuclei stained with DAPI (blue), EFNA4 is detected with anti-EFNA4 (green).
